# Transcriptomic profiles of *Plasmodium falciparum* and *Plasmodium vivax*-infected individuals in Indonesia

**DOI:** 10.1101/2021.01.07.425684

**Authors:** Katalina Bobowik, Din Syafruddin, Chelzie Crenna Darusallam, Herawati Sudoyo, Christine Wells, Irene Gallego Romero

## Abstract

Malaria is one of the leading causes of illness and death globally. The vast majority of transcriptomic studies of the impact of malaria on human hosts have been conducted on populations of African ancestry suffering from *Plasmodium falciparum* infection, making it unclear whether biological responses observed in these studies can be generalised to other populations. Here, we perform differential expression analysis between healthy controls and malaria-infected patients within Indonesia, a country of over 260 million people which has substantial morbidity due to endemic malaria. We find that in samples infected with *P. falciparum* and *P. vivax*, there is an upregulation of genes involved in inflammation, the immediate early immune response, translation, and apoptosis. When comparing these findings to transcriptomic studies conducted in Africa (on *P. falciparum*) and South America (on *P. vivax*), we find that many pathways are shared. This is particularly apparent for receptor recognition and inflammation-related genes in *P. falciparum* and innate immune and chemokine-related genes in *P. vivax* infection. However, we also find that many genes are unique to the Indonesian population, particularly those involved in RNA processing, splicing, and cell surface receptor genes. This study provides a more comprehensive view of malaria infection outside of Africa and contributes to a better characterisation of malaria pathogenesis within humans across a range of genetic architectures.

## Background

Malaria, one of the leading causes of illness and death globally [1], has exerted some of the strongest selective pressures on the human genome [2], but little is known about how the human transcriptional response to malaria varies across populations. Transcriptomic studies conducted on the host response to *Plasmodium* infection have been a critical tool in attempting to explain the functional mechanisms behind malaria pathogenesis, yet the vast majority of transcriptomic studies on the human host have been conducted on populations within Africa [3–5]. It is clear that human adaptation to malaria has occurred locally multiple times [2]. For example, Southeast Asian ovalocytosis, caused by a deletion in the *SLC4A1* gene, confers a broad-spectrum protection against multiple malaria parasites and is found in high frequencies throughout Island Southeast Asia and the Malay Peninsula [6] while HbS, or the sickle haemoglobin allele, is found most commonly in Africa and can result in a 10-fold lower risk of severe malaria [7, 8]. Therefore, there is a crucial need to understand how the host response varies across different ethnic backgrounds, as these differences can impact disease progression and allow for better diagnosis and therapeutics.

Despite being the second-most affected malaria region in the world [9] and hosting significant levels of human genetic diversity [10], Island Southeast Asia—a region which includes the countries of Brunei, Indonesia, Papua New Guinea, the Philippines, and East Timor, as well as smaller island nations—is nearly unrepresented in transcriptomic studies analysing the human host response to malaria. To our knowledge, only one transcriptomic study analysing human gene expression has been conducted on malaria patients within Island Southeast Asia [11], focusing on understanding the relationship between gene expression in humans and *P. falciparum*, as well as how gene expression correlates with clinical data such as malaria severity, age, and body temperature.

However, there is still a lack of understanding of how *Plasmodium*-infected individuals respond to infection compared to healthy controls. Furthermore, *P. vivax*, which differs in its biology to *P. falciparum*, is also a common cause of infection in Island Southeast Asia and the host response to it is even more poorly understood. In order to address these gaps, here we analyse the transcriptional profile of malaria-infected individuals within Island Southeast Asia. Combining previously collected whole blood RNA-seq data from healthy Indonesian individuals [12], with the dataset generated by Yamagishi et al. [11], we analyse which pathways and genes are differentially regulated between patients with simple forms of malaria (uncomplicated and asymptomatic) and healthy controls. Given the different genetic backdrop and strains of *Plasmodium* species within Island Southeast Asia, this study aims to characterise the transcriptomic profile of infection in this region, and determine whether it differs when compared to previous studies—predominantly within the continent of Africa.

## Methods

### Samples

Samples were collated from the only two whole-blood RNA-seq datasets containing malaria-infected and control samples from Indonesia we could find [11, 12]. We combined samples from these two different datasets for two main reasons: First, both datasets are unbalanced between cases and controls, and if analysed separately this would lead to an overrepresentation of one group (e.g., case or control) over the other. Secondly, collating data from both studies results in more samples and therefore increased power to detect differentially expressed (DE) genes.

The first dataset consists of 100 base-pair, paired-end data from whole blood from 116 male individuals (plus 6 technical replicates) collected by members of the Eijkman Institute on the Indonesian islands of Sumba, Mentawai, and on the Indonesian side of New Guinea Island (raw data are available from the European Genome-phenome Archive study EGAS00001003671; as described in [12]), hereafter referred to as the Natri dataset. All collections followed protocols for the protection of human subjects established by institutional review boards at the Eijkman Institute (EIREC #90 and #126); the analyses in this publication were additionally approved by University of Melbourne’s Human Ethics Advisory Group (1851585.1).

The second dataset consists of 36 base-pair, single-end data from whole blood collected from individuals within Sulawesi, Indonesia [11], hereafter referred to as the Yamagishi dataset. Raw sequencing reads for cases (n = 122) and controls (n = 25) were downloaded from SRA studies DRP000987 and DRP001953, respectively.

### RNA sequencing data processing

RNA-seq reads from both studies went through an initial sample quality analysis using FastQC V. 0.11.5 [13]. Leading and trailing bases below a Phred quality score of 20 were removed using Trimmomatic v. 0.36 [14]. RNA-seq reads were then aligned to the human genome (GrCh38, Ensembl release 90: August 2017) with STAR v. 2.5.3a [15] using a two-pass alignment and default parameters. The maximum number of mismatches was set at 10 for the Natri dataset and 3 for the Yamagishi dataset to account for differences in read lengths, which resulted in a mean of 29 million and 17 million uniquely-mapped read pairs for the Natri and Yamagishi datasets, respectively (Supplementary Table 1). All reads which did not map to the human genome were then remapped with STAR [15] to a combined *P. falciparum* (PF3D7) and *P. vivax* (PVP01) genome obtained from PlasmoDB (release 36) [16]. The following additional flags were added to account for small genome size: genomeSAindexNbases = 11, alignIntronMax = 35,000 and alignMatesGapMax = 35,000. We assigned reads to the genome with the highest mapping quality; when reads mapped to both genomes with the same mapping quality, they were assigned to both species.

We performed human read quantification with featureCounts v. 1.5.3 [17] and used a filtered GTF file which only included GENCODE basic (release 27; August 2017) annotation with transcript support levels 1-3. We then used the default settings for single (Yamagishi) and paired end (Natri) reads. On average, we successfully assigned 15 million paired-end reads to each sample from the Natri dataset and 13 million for the Yamagishi dataset (Supplementary Table 1).

For the Natri dataset, library preparation was performed using the Globin-Zero Gold rRNA Removal Kit (Illumina), however a similar depletion step was not performed during the library preparation stage for the Yamagishi dataset. We therefore bioinformatically removed all globin genes from the study (*HBA1, HBA2, HBB, MB, CYGB*, and *NGB*) which resulted in a more continuous distribution of the total mRNA read pool (Supplementary Figure 1). The minimum library size was then set to 9 million reads post globin removal, which was the minimum number of reads in the Natri dataset. This reduced the total sample size of the Yamagishi dataset from 147 samples to 95.

### Deconvolution analysis

In order to control for heterogeneity within blood cell type proportions and parasite developmental stages, we performed blood and parasite stage deconvolution as follows: For whole blood, we used DeconCell v. 0.1.0 [18] to estimate the proportion of granulocytes, B cells (CD19+), CD4+ T cells, CD8+ T cells, natural killer (NK) cells, and monocytes (range of deconvoluted proportions: 82.5%-99.9% for the Yamagishi data and 89.4%-100% for the Natri data, the latter numbers were taken from [12]). Estimates of the predicted cell proportions are available in Supplementary Table 2. To assign parasite life cycle stage, we used a file from Aunin et al. [19] which contains single cell CPM-normalised gene expression measurements calculated across 12 different parasite stages taken from *Plasmodium berghei*. Since this study focuses on the human whole blood transcriptome, only Plasmodium stages within the human blood cell stage were used (male and female gametocyte, trophozoite, ring, merozoite, and schizont stages). *P. vivax* and *P. falciparum* gene names were converted to one-to-one orthologous *P. berghei* gene names using an orthologous gene file obtained from the Malaria Cell Atlas [20]. We then removed all samples which had no reads mapping to either *P. falciparum* or *P. vivax* and uploaded the CPM-normalised counts to CIBERSORT [21] using 100 permutations and disabling quantile normalisation (as recommended on the CIBERSORT page). This was performed separately for reads mapping to *P. vivax* and *P. falciparum*, and reads mapping equally well to either species were included in both files. The output from CIBERSORT can be viewed in Supplementary Table 3.

### Sex classification

Due to missing sex information within the Yamagishi dataset, we assigned sex based on expression of marker genes and clustering of reads mapping to the X and Y chromosomes. The *XIST* gene was used as a female expression marker [22] and *RPS4Y1, EIF1AY, DDX3Y*, and *KDM5D* were used as male expression markers [23]. This resulted in 8 unlabelled samples mapping to the female group and 5 unlabelled mapped to the male group.

### Defining malaria samples and species identification

The malaria infection status of samples was assessed as follows: in the Natri dataset, four samples showed evidence of *Plasmodium* infection through nested PCR—a method considered to be the “gold standard” for *Plasmodium* detection [24]. We then quantified the fraction of reads across these four individuals that mapped to the combined *Plasmodium* genomes (total number of mapped reads/total number of reads in the starting library), and used this as a threshold for classifying samples as having malaria. All samples which passed this initial threshold (0.04%) also had to have reads mapping to a minimum number of genes, where the PCR-confirmed sample with the lowest number of genes (n = 807) was again used as the baseline. Using this criteria, six samples in this dataset were designated as having malaria (For the full reassignment of samples, see Supplementary Table 4).

For the Yamagishi dataset, the authors originally confirmed malaria status through either a rapid malaria paper test or by a smear test. Both of these tests are well documented as commonly having false positives and false negatives [25–27]. Therefore, we classified malaria status by the following: since control samples also had a low number of reads mapping to *Plasmodium*, we took the control sample with the highest fraction of reads (0.32%) and number of genes (n = 506) mapping to *Plasmodium*. Any sample that did not exceed both of these thresholds was considered to be healthy. Using this criteria, one sample appeared to be healthy and was removed from the analysis, as it could potentially reflect the expression profile of another illness (all sick individuals from the Yamagishi study were collected within a hospital and self-proclaimed sick; For the full reassignment of samples, see Supplementary Table 4.)

Finally, to assess whether individuals were infected with *P. falciparum* or *P. vivax*, we took into account the number of reads mapping to either genome for each individual (Supplementary Table 5), as well as clustering by PCA. Samples were first classified as infected by either *P. falciparum* or *P. vivax* if over 50% of *Plasmodium* reads mapped to that species. PCA using the *Plasmodium* data, (Supplementary Figure 2) confirmed this assignment, with samples assigned to either species seen to form two distinct clusters, and samples classified as healthy (as above) forming a third. The exception to this was within three samples belonging to the Yamagishi data (DRR006381, DRR006392, and DRR006431). Although these three samples had more reads mapping to *P. vivax*, they were seen to cluster with *P. falciparum* samples in PCA, as well as hierarchical clustering of the *Plasmodium* reads (Supplementary Figure 2). These samples were therefore classified as having *P. falciparum* infection.

### Differential expression analysis

We set filtering parameters to only retain genes that were expressed at a CPM ≥ 1 in at least half of individuals from designated case or control groups. Using these parameters, we kept 12,895 genes from the Yamagishi dataset and 12,743 from the Natri dataset. We then merged both datasets and normalised the library using TMM normalisation [28], resulting in a matrix of 11,735 genes.

To account for variables affecting gene expression levels, we used an ANOVA to test for the association between the first three principal components of the data which captured over 70% of the variation and all known covariates. Age, which is associated with malaria severity [29], was not available for nine individuals from the Yamagishi study. We therefor removed these nine samples from the dataset, resulting in a total of 201 samples (plus 6 replicates). The breakdown of samples can be viewed in Table 1.

**Table 1.** Number of samples by sex, study, and location after filtering.

| Island | Study | Sex | Case (Control) | Samples |
| --- | --- | --- | --- | --- |
| Mentawai | Natri | M | 0 (48) | 48 |
| Sumba | Natri | M | 1 (48) | 49 |
| New Guinea Island | Natri | M | 5 (14) | 19 |
| Sulawesi | Yamagishi | F | 20 (10) | 30 |
|  | Yamagishi | M | 43 (12) | 55 |
| **Total** |  |  |  | **201** |

After accounting for variables affecting gene expression, the final linear model used to test for differential expression was the following:

Expression ~ species + island + age + sex + falciparum stage proportion (female, male, merozoite, ring, schizont, trophozoite) + vivax stage proportion (female, male, merozoite, ring, schizont, trophozoite) + blood cell type proportion (CD8T, CD4T, NK, B cells, monocytes and granulocytes)

As this model contains a large number of parameters (22), we wanted to ensure that we were not overfitting it to the data. Therefore, we also tested 3 additional models with reduced parameters using Akaike information criterion (AIC [30]) from the selectModel function in limma v. 3.42.2 [31]: removing blood deconvolution estimates, removing *Plasmodium* stage estimates, and removing blood and *Plasmodium* stage estimates. We found that the full model including all parameters greatly outperformed all other models (6,803 genes were associated with the full model being the best fit, followed by 4,341 genes in favour of a model without *Plasmodium* stage estimates, 324 genes for a model with only species, island, age, and sex information, and 267 genes for a model without blood proportion estimates). For a full breakdown of the estimated proportion of variance each covariate contributed to expression levels, see Supplementary Figure 3.

Differential expression analysis was then conducted using limma with high sample variability removed using the Voom function [32]. We used the technical replicates from the Natri dataset to capture the random individual effect using the limma function duplicateCorrelation [33]. An absolute log_2_(FC) threshold of 1 and an FDR-corrected (Benjamini-Hochberg [34]) p-value of 0.05 was used in order to identify differentially expressed genes.

### Gene set analysis

To explore enriched gene sets, we used ClueGo v. 3.8.0 [35], a Cytoscape plugin to decipher GO networks and pathways. The following parameters were used for the GO analysis: Statistical Test Used = Right-sided hypergeometric test for enrichment; Correction Method = Benjamini-Hochberg; Minimum Number of Genes = 4; Minimum Percentage of Genes = 2%. To analyse biological pathways, we used Reactome, a curated database of biological pathways [36, 37].

### Comparison of the Indonesian gene sets to other gene sets

In order to assess whether the Indonesian gene sets were similar to other differential expression analyses, we compared the *P. falciparum* and *P. vivax* gene sets to three other publicly-available studies conducted on uncomplicated malaria patients versus healthy controls. The first study was performed on febrile adults infected with *P. falciparum* within Mali, an endemic malaria region [38]. Since samples from children are also present in this study, we also collected data from *P. falciparum*-infected children under the age of 10 in Benin [4]. For the *P. vivax* data, we retrieved data from a controlled human malaria infection (CHMI) study conducted on malaria exposed and malaria naive individuals within Colombia [39]. In order to allow for differences in study design, we did not use an absolute log_2_ FC threshold cut-off in comparison to these studies and only filtered for genes with an FDR-corrected p-value less than 0.05.

## Results

### Exploration and control of covariates influencing expression levels

In order to provide a more comprehensive understanding of malaria pathogenesis in Island Southeast Asia, we have combined data from two RNA-seq studies from malaria-infected and control samples from the Indonesian archipelago. We reanalysed whole blood transcriptomes obtained from 69 Indonesian individuals with simple forms of malaria (asymptomatic and uncomplicated), infected with *P. falciparum* and *P. vivax* and compared them to 132 healthy controls. After excluding samples with low library sizes and removing lowly-expressed genes, this resulted in a median library size of ≈15 million for human reads (range:≈ 9 × 10^6^ − 3.2 × 10^7^) and ≈ 600,000 for reads mapping to the *P. falciparum* and *P. vivax* genomes (range:≈ 8 × 10^3^ − 9 × 10^6^).

To explore which variables might be driving variation within the data, we performed principal component analysis (PCA) on the human log_2_-normalised CPM values. As expected, study had the strongest association with expression levels (Supplementary Figure 4; Supplementary Table 6). Island also had a highly significant association with expression levels, however this was driven predominantly by study, i.e., the island of Sulawesi (Yamagishi) versus the other three islands: Mentawai, Sumba, and the Indonesian half of New Guinea Island (Natri; Supplementary Figure 4). We also observed that leukocyte proportions, which are influenced by infection, were all significantly associated with expression levels (Supplementary Table 6).

As parasite stage can influence gene expression [3, 40, 41], we deconvoluted the *Plasmodium* read data through a reference signature gene file from Aunin et al. [19]. We found that for both the *P. falciparum* and *P. vivax* data, the most predominant stages were early stage circulating forms of the parasite (merozoites and ring stages, Figure 1, A and B). This aligns with previous research from the Malaria Cell Atlas which has shown that in *P. falciparum* bulk data, the majority of the cell population is within early stage forms. For *P. falciparum*, the merozoite stage—the stage in which the parasite is directly exposed to the host immune system [42]—was most prevalent.

**Fig 1.**
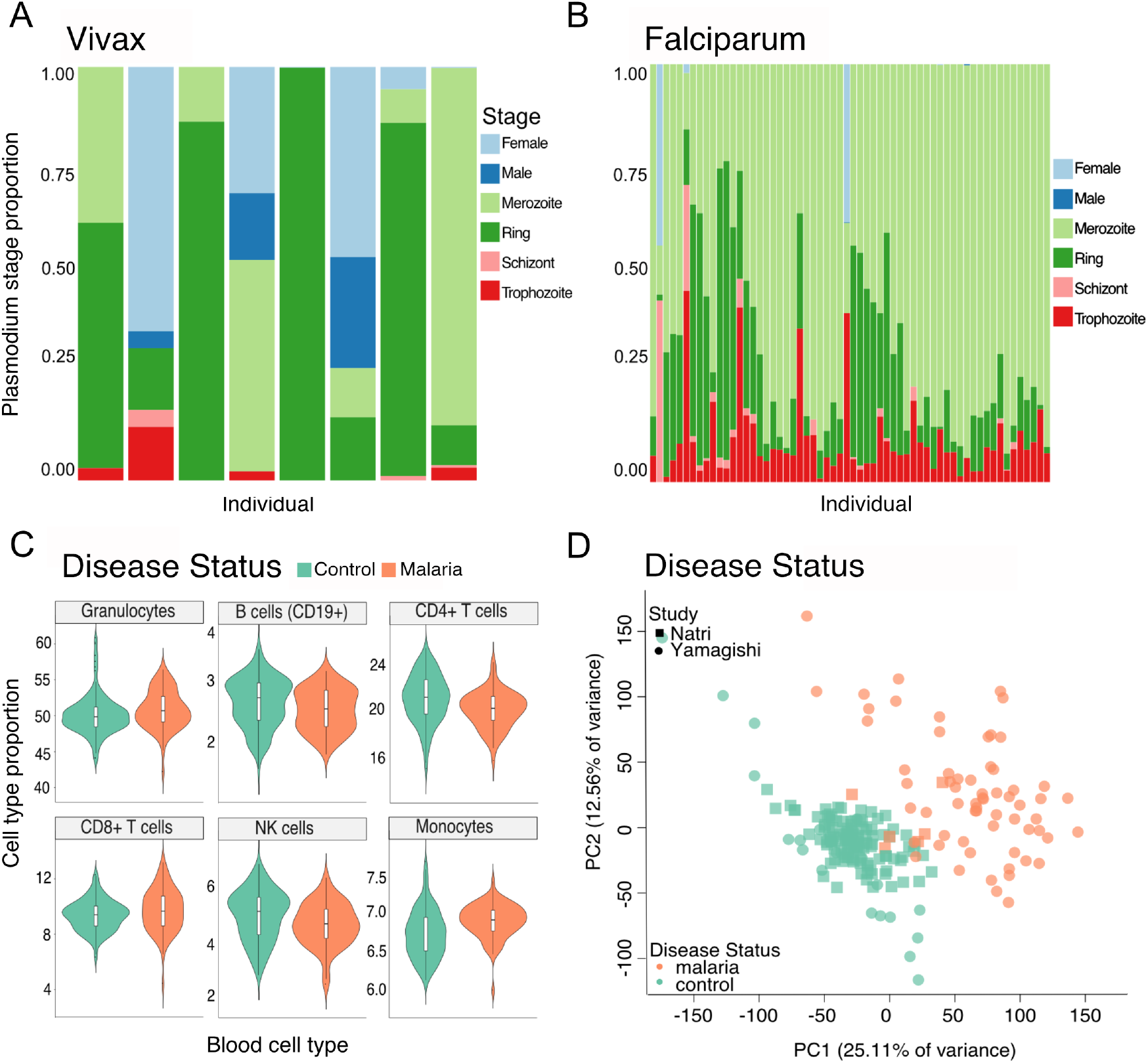
Covariates influencing expression levels. A) Estimated *Plasmodium* stages within *P. vivax* and B) *P. falciparum*. C) Violin plots showing the proportion of blood cell types for each leukocyte within control and malaria-infected individuals. D) PCA analysis of disease status after correcting for batch effects within control (green) and malaria (orange) individuals.

We observed a significant difference between control and case samples in levels of B cells, CD4T cells, NK cells, and monocytes (BH-corrected ANOVA p = 0.018, 5.7×10^−4^, 0.007, and 1.4×10^−4^, respectively; Figure 1, C). Out of these cell types, control samples had the higher cell type proportion with the exception of monocytes, which were higher in individuals with malaria. Tukey’s HSD post-hoc test on CLR-transformed data resulted in significant levels for CD4T cells, CD8T cells, NK cells, and monocytes (BH-corrected ANOVA p = 0.005, 0.018, 0.017, and 5.1 × 10^−4^, respectively), with control samples having higher levels of CD4T cells and NK cells, and cases having higher levels of CD8T cells and monocytes (Supplementary Table 2). As noted in previous studies, this increase in monocytes within cases may indicate a high pro-inflammatory immune response [43] and may be protective in order to decrease parasite burden [44, 45].

After adjusting for island, sex, age, stage proportion, and blood cell type proportion, we found that disease status captured most of the remaining variation in the data, followed by *Plasmodium* species (Supplementary Table 6). Grouping by PCA showed two distinct clusters for control and case samples, with samples from both studies clustering in either group (Figure 1, D). Although we observed clustering between both control and case groups, we observed more variance in the case group than in the control group. We attribute some of this observation to differences in disease severity between malaria-infected individuals; Yamagishi et al. [11] reported a linear association between malaria severity and *Plasmodium*-derived read abundance within their samples. Furthermore, as the majority of malaria samples are drawn from the Yamagishi data, which is single-ended and has a shorter read length than the Natri data, the higher variance within the malaria samples could be due to more variation in alignment confidence, particularly across highly variable regions of the genome [46]. This is supported by the fact that the Natri dataset had a much higher proportion of reads that mapped uniquely to the human genome (median = 87%) compared to the Yamagishi dataset (median = 58%; Supplementary Table 1).

### Enriched genes in *P. falciparum* and *P. vivax* are involved in the immune response, translation, and apoptosis

As we are interested in the biological response to malaria within the Indonesian population, we evaluated the magnitude and significance of differentially expressed genes between cases and controls and separated this into the different infecting *Plasmodium* species: *P. falciparum* and *P. vivax*. Table 2 summarises the results of differential gene expression testing in both group comparisons. At a log_2_ FC threshold of 1, a total of 1,003 genes were differentially expressed in either patients infected with *P. vivax* or patients infected with *P. falciparum* (2,559 genes without any threshold). Of these, 150 genes (373 genes without any threshold) were shared, 639 were unique to *P. falciparum* (1,811 genes without any threshold), and 214 were unique to *P. vivax* (375 genes without any threshold).

**Table 2.** Number of differentially expressed genes in *P. falciparum* and *P. Vivax* -infected individuals compared to healthy controls at an adjusted p-value (BH) of 0.05 and a log_2_ FC threshold of 1 (no log_2_ FC in parentheses).

|  | **Falciparum vs Control** | **Vivax vs Control** |
| --- | --- | --- |
| Down | 345 (1,083) | 112 (293) |
| NotSig | 10,946 (9,551) | 11,371 (10,897) |
| Up | 444 (1,101) | 252 (455) |

Many of the genes which were DE in both *P. falciparum* and *P. vivax* were part of Reactome pathways related to signalling by interleukins, translation, nonsense mediated decay, and diseases such as influenza (Supplementary Table 7). As identified by Reactome pathway analysis, these shared genes contribute to pathways involved in malaria infection, some of which include regulation of apoptosis [47], inflammation and the immune response [48], and cellular response to environmental stress [49]. For a full list of shared genes, see Supplementary Table 8.

Of all *Plasmodium* species capable of infecting humans, *P. falciparum* is responsible for the most severe form of malaria [50], and therefore we investigated the effects of this species versus healthy controls separately. Some of the most highly upregulated genes in *P. falciparum*-infected individuals included immediate early response genes such as *FOS* [5] and *DUSP1* [51], as well as genes involved in apoptosis (some of the most highly upregulated genes being *G0S2* and *BCL2A1*). We also observed a strong signal from inflammation-related genes; *SOCS3* and *IFNG*, two major regulators of infection and inflammation [52, 53], were significantly upregulated in *P. falciparum* patients. Furthermore, inflammatory chemokines, which are well-documented to have an association with malaria [54], were both significantly up and down regulated (Figure 2). For example, expression of *CXCL8* was upregulated, as noted in other studies [54–56], while *CCR3* and *CCR4* were downregulated, an effect which has previously been observed [38].

**Fig 2.**
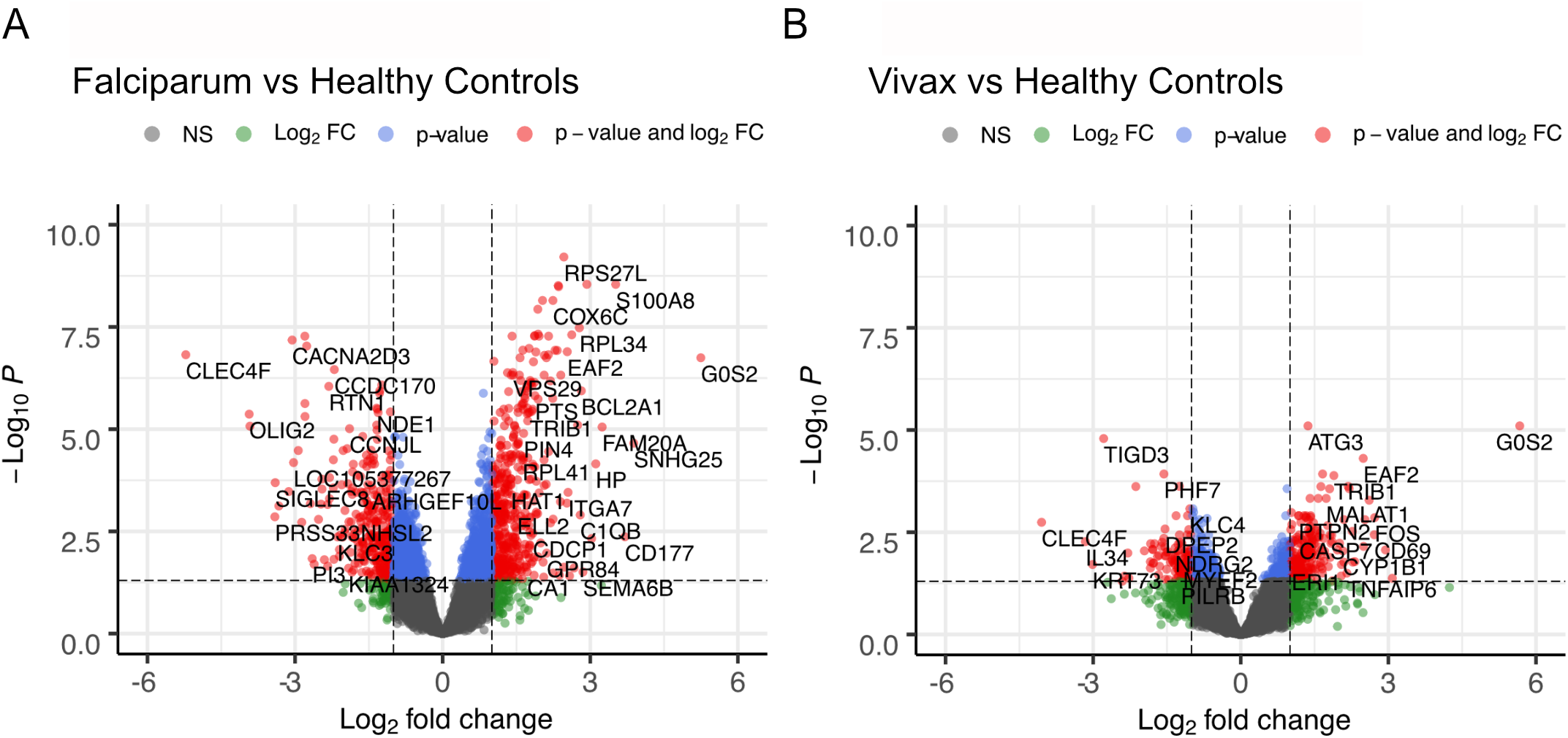
Genes and pathways enriched in *P. falciparum* and *P. vivax*-infected individuals. A) Volcano plot of significant genes (indicated in red) at an absolute log_2_ FC of 1 and a p-value cut-off of 0.05 for *P. falciparum* and B) *P. vivax*-infected individuals.

Some of the most highly upregulated genes in our data were genes encoding ribosomal proteins, as well as genes involved in translation pathways (Supplementary Table 9). In addition, some of the most significantly-enriched Reactome pathways in *P. falciparum* patients were those involved in translation and nonsense mediated decay (Supplementary Table 7), and GO analyses were enriched in the regulation of apoptotic processes and translation (Supplementary Table 10). Since ribosomal protein genes are from large gene families [57], we tested whether this signal could be driven by differences between cases and controls in read length (the majority of reads from cases being 36 bp long, while those from controls being 100 bp long in most cases). We found that when we stratified cases and controls by read length, mean expression of significant DE genes involved in translation (beginning with RPL, MRPL or RPS; n = 85) amongst sample classes were still significantly different (Tukey’s HSD on 36 bp cases versus 100 bp controls ANOVA p = 3.4 × 10^−8^; Supplementary Figure 6), suggesting a biological response rather than sequencing artefact.

Due to differences in biology to *P. falciparum* [58, 59], we also analysed the effects of infection with *P. vivax* compared to healthy controls. We found that Reactome pathways were significantly enriched in pathways related to interleukin signalling (Supplementary Table 7), while GO terms were enriched in cellular response to stress, regulation of metabolic processes, response to cytokines, translation, and apoptotic processes (Supplementary Table 10).

As above, some of the most highly enriched genes in *P. vivax* patients were those involved in the immediate early immune response, inflammatory response, and translation (Supplementary Table 9). For all shared genes, we also found changes due to *P. vivax* infection to have the same direction of effect and similar log_2_ FC intensities to those observed in *P. falciparum* patients (Supplementary Figure 5). However, some differences with the *P. falciparum* response were also apparent: We found that *P. falciparum*-infected individuals had a much stronger signal in the defense and inflammatory response (Supplementary Table 9, Supplementary Table 10), as shown by an upregulation of chemokines and *IFNG*, both of which were absent in *P. vivax*-infected individuals. *ICAM1* and the complement component genes *C1QA* and *C1QB*, all of which have previously been associated with malaria severity [60, 61], were also upregulated exclusively in *P. falciparum*-infected patients. Finally, we found that *CD69*, a marker of recent T-cell activation which has previously been associated with less severe forms of malaria [62], was one of the most highly upregulated genes in *P. vivax*-infected individuals (BH-adjusted p = 1.9 × 10^−5^), but not significantly differentially expressed in *P. falciparum*-infected patients (BH-adjusted p = 0.052).

### Comparing the Indonesian response to malaria infection to other populations

In order to identify any possible Indonesian-specific component of this response, we compared genes differentially expressed in Indonesia in response to *P. falciparum* and *P. vivax* infection to four other publicly-available datasets of differentially expressed genes, taken from three studies conducted on uncomplicated malaria patients versus healthy controls [4, 38, 39]. The first two datasets are from *P. falciparum*-infected individuals who have either previously had malaria exposure (malaria-exposed individuals), collected from individuals in Mali [38], or individuals who were never exposed to malaria (malaria-naive), taken from children in Benin [4]. The other two datasets are from *P. vivax*-infected individuals from Colombia, collected from both individuals living in malaria-endemic regions in Buenaventura (malaria-exposed), as well as individuals from Cali with no previous malaria exposure (malaria-naive) [39]. In all cases, the available data is the list of differentially expressed genes between patients and healthy controls, and we tested for overlap between lists without thresholding on log_2_ FC values.

For comparisons to *P. falciparum*-exposed individuals from Mali, we began with a list of 2,775 genes found to be differentially expressed by the authors (p = 0.05; number of individuals = 3; RNA-seq measurements). Of these, 541 genes were differentially expressed in our own data out of 2,569 genes (21.06%) which were present in both datasets. For comparisons to *P. falciparum*-exposed individuals from Benin, we took a list of 2,351 genes found to be differentially expressed by the authors (p = 0.05; number of individuals = 94; microarray measurements). Of these, 425 genes were differentially expressed in our own data out of 2,098 genes (20.26%) which were present in both datasets (Figure 3). Many of these genes are involved in apoptosis, inflammation, and the innate immune system. Furthermore, we found many of these genes shared the same direction of effect and had a similar magnitude of expression. This is particularly true in comparison to the population within Mali, where nearly all genes shared the same direction of effect and had similar trends in fold change (R = 0.82).

**Fig 3.**
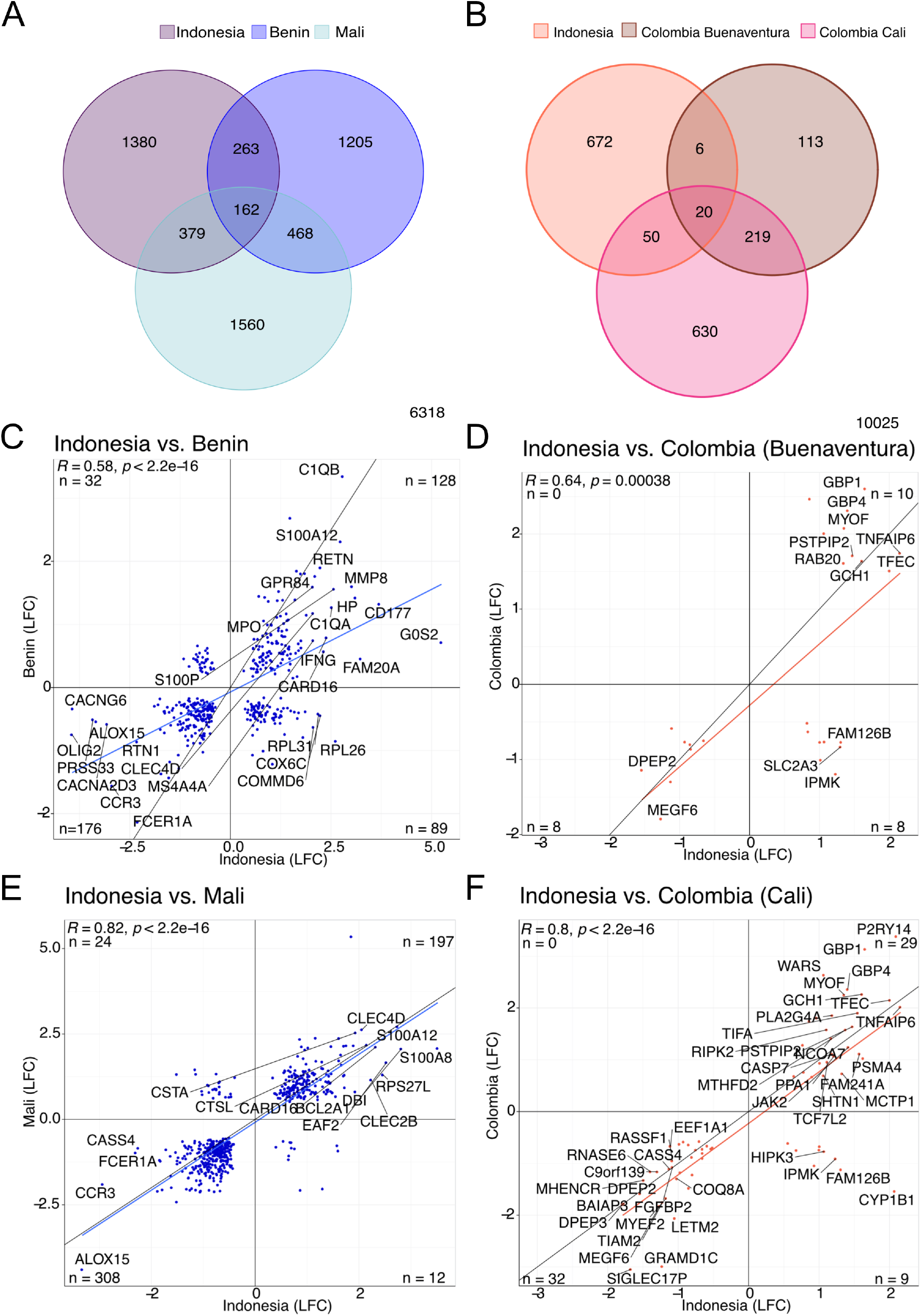
Comparisons between Plasmodium-infected patients in Indonesia and other malaria-endemic countries. A) Total number of differentially expressed genes between *P. falciparum*-infected individuals and healthy controls in Indonesia, Benin, and Mali (FDR ≤ 0.05 and no absolute log_2_ FC). B) Total number of differentially expressed genes between *P. vivax*-infected individuals and healthy controls in Indonesia, Cali, Colombia, and Buenaventura, Colombia (FDR ≤ 0.05 and no absolute log_2_ FC). C-F) Comparison of enriched genes (FDR ≤ 0.05 and no absolute log_2_ FC) within Indonesian populations and populations outside of Island Southeast Asia [4, 38, 39]. Panels C and E are enriched genes for *P. falciparum* while panels D and F are for genes enriched in *P. vivax*.

For comparisons to *P. vivax*-exposed individuals from Colombia, Cali who are malaria-naive, we took a list of 1,072 genes found to be differentially expressed by the authors (p = 0.05; number of individuals = 6; RNA-seq measurements). Of these, 70 genes were differentially expressed in our own data out of 919 genes (7.62%) which were present in both datasets. For comparisons to *P. vivax*-exposed individuals from Colombia, Buenaventura who had prior malaria exposure, we took a list of 400 genes found to be differentially expressed by the authors (p = 0.05; number of individuals = 6; RNA-seq measurements). Of these, 26 genes were differentially expressed in our own data out of 358 genes (7.26%) which were present in both datasets (Figure 3). Many of these genes are involved in innate immunity, such as the interferon-induced genes *GBP1* and *GBP4*, as well as *JAK2*. Other shared genes include apoptosis-related genes and transport genes (Figure 3). We found many more genes were shared between populations from Cali, Colombia (the malaria-naive population) than populations from Buenaventura, Colombia (the malaria-exposed population), as well as a stronger correlation in log_2_ FC magnitude and direction (R = 0.8).

In our comparison of responses to infection with *P. vivax* and *P. falciparum*, we found that many differentially expressed genes were shared across populations in response to infection by either species. In order to test, on a broader scale, which genes are shared between not only species but also different global regions, we compared all significant genes (FDR ≤ 0.05 with no log_2_ FC threshold) within the Indonesian population to the African (combining the differentially expressed genes within Benin and Mali for a total list of 4,037 genes) and South American populations (composed of the differentially expressed genes within the Colombian populations of Cali and Buenaventura, again combined, 1,038 genes). We found that many genes are unique to the Indonesian populations (unique to *P. falciparum* infection versus all non-Indonesian datasets = 1,072; unique to *P. vivax* infection versus all non-Indonesian datasets = 141; common across *P. vivax* and *P. falciparum* infection but absent from all non-Indonesian datasets = 177), although we observed a substantial overlap across the populations in our study (Figure 4). Indonesian populations infected with either *P. falciparum* or *P. vivax* had the highest similarity with populations from Africa. Furthermore, 29 genes were shared between all species and all populations, suggesting that despite *Plasmodium* species and genetic background, some host genes are integral during *Plasmodium* infection (see Supplementary Table 8).

**Fig 4.**
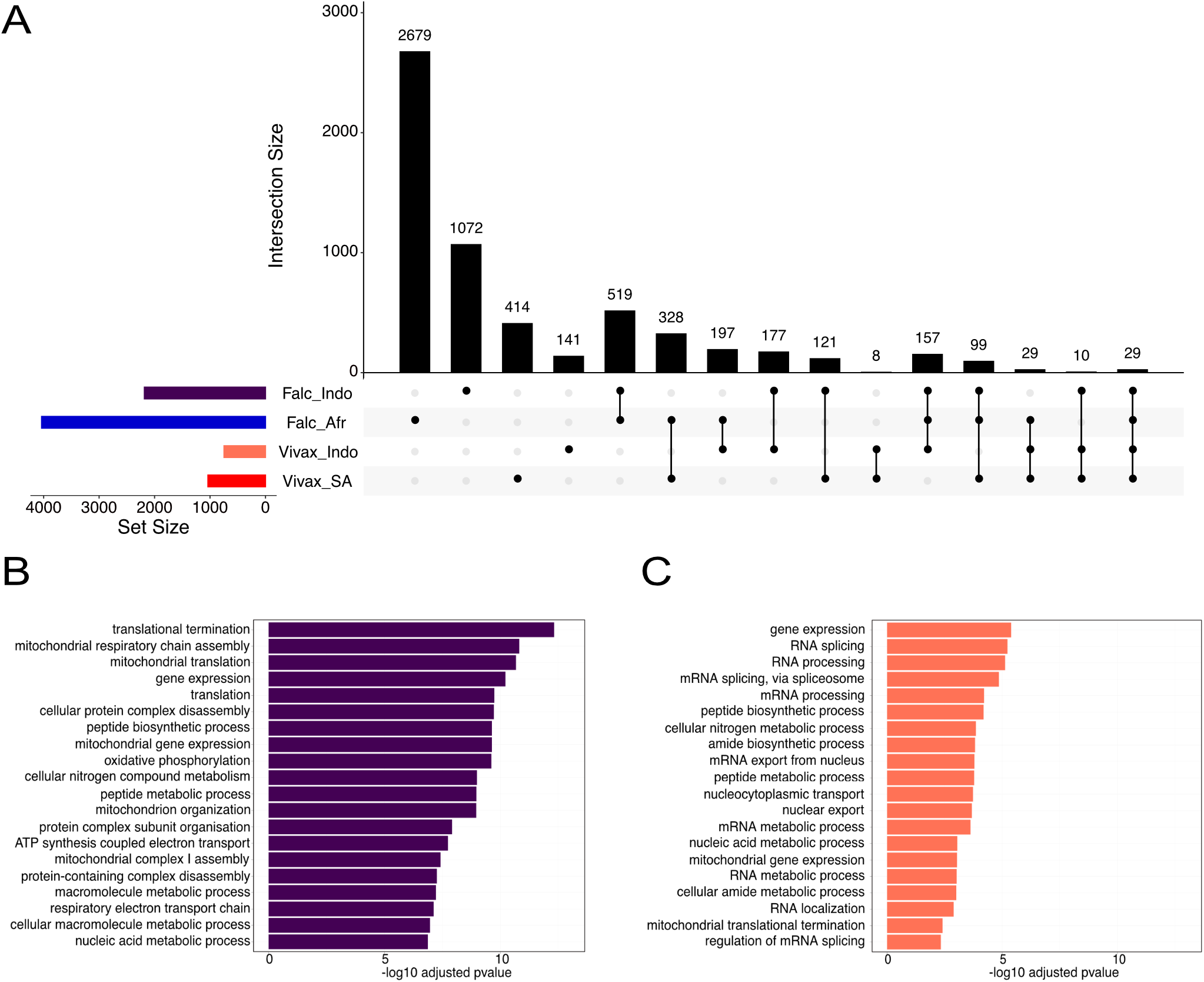
Shared and Indonesian-specific genes within *P. vivax* and *P. falciparum*-infected patients. A) Shared and unique genes across multiple populations. *P. falciparum* and *P. Vivax*-infected individuals within Indonesia compared to *P. falciparum* and *P. vivax*-infected individuals within South America (Colombia) and Africa (Mali and Benin). The number of shared genes for each comparison can be viewed on the top of each bar. B) Gene ontology enrichment terms (BP) for unique genes found within Indonesian individuals infected with *P. falciparum*. The top 20 terms by adjusted p-value (BH) are shown. C) Gene ontology enrichment terms (BP) for unique genes found within Indonesian individuals infected with *P. vivax*. The top 20 terms by adjusted p-value (BH) are shown.

Finally, we wanted to investigate those genes whose upregulation appears unique to malaria-infected individuals within Indonesia, (n = 1,213; Figure 4). Although differences in study design and technology between studies affects our ability to definitively attribute these genes as being Indonesian-specific, exploring these genes allows for an initial investigation of potential population-specific pathways. For individuals infected with *P. falciparum*, we found many of these genes were enriched in GO pathways related to RNA processing and splicing (Figure 4; for a full list of enriched GO terms, see Supplementary Table 10), as well as Reactome pathways involved in mitochondrial translation, respiratory electron transport, and activation of ATR in response to replication stress (Supplementary Table 7). For genes specific to *P. vivax*, we found the majority of genes were also enriched in GO terms and Reactome pathways involved in RNA processing and splicing, as well as translation and protein transport (Figure 4; Supplementary Table 7; Supplementary Table 10). Indeed, genes such as Sm-like protein genes and SR protein genes, which are crucial to RNA splicing, were upregulated within *P. falciparum* and *P. vivax*-infected patients. Other notable genes include surface expressed protein genes such as *CLEC4F* and *SIGLEC8*, which had some of the largest log_2_ FC values. (The full list of genes unique to individuals infected with *P. falciparum* and *P. vivax* can be found in Supplementary Table 9.)

## Discussion

Our understanding of the human immune response to malaria infection remains in its infancy. Here, we have investigated the transcriptional response to simple forms of malaria within a large group of Indonesian individuals and find both conserved and unique components to it, relative to other human groups. We found a total of 789 genes differentially expressed in response to infection by *P. falciparum* and 364 in response to *P. Vivax*, within the range of previous findings.

The innate immune system plays an important role against pathogens by eliciting physical and chemical barriers to protect the host [63], and we found that this was one of the strongest signals observed in our data. Inflammation, a pathway which mediates parasite clearance [64], was evident by the upregulation of *IRAK3, S100A8*, and *SOCS3*, all known inflammatory markers and mediators [52, 65, 66]. This finding corroborates earlier research which has shown that the inflammatory response can be triggered by intracellular parasites which destroy infected cells [48]. Other innate immune signatures included the upregulation of *FOS*, a gene implicated in the immediate early immune response and immune modulation [67], as well as the upregulation of genes involved in the complement system, a pathway which plays an important role in the inhibition of merozoite invasion [68].

Beyond innate immune responses, we also observed expression of genes involved in apoptosis. Apoptosis is a mechanism which removes damaged cells from the body [69] and is an observed effect during malaria infection [70–72]. Although the role of apoptosis in malaria infection has not been fully elucidated [73], some studies have proposed that apoptosis occurs in response to haemozoin, a waste product of haemoglobin digestion by Plasmodium [71, 72, 74, 75]. Indeed, the upregulation of apoptosis-related pathways and genes, particularly *G0S2* which had the highest log_2_ FC value in both *P. vivax* and *P. falciparum*, could therefore be a response to haemozoin and oxidative stress.

In order to explore similarities in the regulatory mechanisms of malaria infection, we compared differentially expressed genes from *P. falciparum* and *P. vivax*-infected Indonesian individuals to three other transcriptomic studies conducted on individuals with uncomplicated forms of malaria and healthy controls from Africa [4, 38] and South America [39], bearing in mind the fact that there are substantial differences in sample size, power and study design between all the data sets we considered. We found that many shared genes in *P. falciparum*-infected individuals were involved in immune-related processes such as receptor recognition genes, chemokine receptors, and inflammation-related genes. In comparisons between *P. vivax*-infected populations, innate immune genes such as *JAK2* and interferon-induced genes, were shared. Malaria is a highly inflammatory disease, and therefore these shared genes may reflect a general inflammatory response.

These shared genes exhibited similar direction of effects across studies (Figure 3), which was particularly apparent for malaria-exposed adults from Mali (R = 0.82). Comparisons to the Benin population had a lower correlation coefficient (R = 0.58) to that of Mali, which may be a reflection of technological differences, as the study was conducted using microarray data in 2012 [4]. We also note that our *P. vivax* patient dataset and the Colombian populations we used for comparison to our *P. vivax* data have small sample sizes (n = 6 and n = 8, respectively), which may explain the lower correlations we observed, especially against malaria-exposed individuals from Buenaventura (R = 0.64; Figure 3).

We also attempted to identify malaria-associated pathways which are unique to Indonesia. We found that many of the enriched genes unique to the Indonesian population were heavily involved in RNA processing and splicing, and that these differences were not due to differences in sequencing length between groups (Supplementary Figure 6). RNA splicing and its impact on immunity has been described previously (for an extensive review, see [76]) and studies have shown that variation in alternative splicing is common across populations from different ethnic backgrounds [77]. No genes known to be implicated in malaria resistance or susceptibility were found within the Indonesian dataset, however we did find that surface expressed protein genes such as *CLEC4F* and *SIGLEC8*, which had some of the largest log_2_ FC values, were unique to the Indonesian dataset. Receptors expressed on immune cells are important mediators of the immune response [78, 79], and previous research has implicated other C-type lectins, such as *CLEC12A* [80] and *CLEC4A* [81], in modulating malaria pathogenesis. This may suggest that the variability in ways that populations differ from one another are still not fully understood and may include variation in genes such as alternative splicing, receptor processing, and binding.

All of the studies we have considered used different approaches to data generation and processing. Despite our best efforts, some unwanted technical and biological variation is likely to remain in the data. However, despite these differences, many of the genes differentially expressed in our Indonesian data after batch correction were also genes with some of the strongest signals in other studies that we compared to. These overlapping genes are involved in antigen clearance, cellular signalling, and inflammation, which are all well-documented pathways involved in malaria infection [82–84]. This may suggest that many crucial immunological pathways during infection are shared across populations, a finding which could be particularly useful for biomarker discovery and diagnostics in the field. While some of the differences we observe between populations may be artefactual, population-specific responses to malaria are well documented and not unexpected [2].

Studying the biological response to infection within the human host across a range of genetic backgrounds is crucial for a more comprehensive understanding of disease, and by extension, the efficacy of vaccines [85], drugs [86], and biomarkers [87]. This is particularly true for malaria, where it has been shown that populations vary in their metabolism of drugs and can suffer from severe reactions to antimalarials [88]. Broadening the repertoire of genes known to be implicated in malaria across populations is essential if we aim to better diagnose and treat this disease. By characterising that response in populations within Indonesia, and situating it within the global context, we have shown that the immune response to malaria infection can have shared and population-specific effects, and that considering these population differences is an essential step in creating more broadly effective malaria therapeutics.

## Author’s contributions

KSB and IGR designed the study. KSB performed analyses and wrote the manuscript with input from all authors. DS performed PCR malaria diagnosis. HS and CCD generated the Indonesian datasets.

## Acknowledgements

We would like to acknowledge all of the study participants who generously consented to genome sequencing in the original study. We would also like to thank Pradiptajati Kusuma (Eijkman Institute) for his valuable comments. KSB was supported by a Melbourne Graduate Research Scholarship.

## Supplementary materials

**Supplementary Figure 1: Globin Gene Removal:** Globin Gene Removal in the Yamagishi dataset.

**Supplementary Figure 2: Species summary:** Principal component analysis of reads mapping to *P. falciparum* and *P. vivax*.

**Supplementary Figure 3: Variance partition:** Violin plots of the percentage of variance explained by all covariates used in the linear model.

**Supplementary Figure 4: PCA of Island:** PCA analysis of island-level variation.

**Supplementary Figure 5: Variance between *P. falciparum* and *P. vivax* genes**

**Supplementary Figure 6: Ribosomal protein genes:** Batch-corrected log_2_ CPM values of ribosomal protein genes, compared between read length and disease status.

**Supplementary Table 1** The number of reads at each filtering stage for both datasets.

**Supplementary Table 2** Blood proportion estimates calculated by DeconCell [18] and BH-corrected ANOVA p-values of Tukey’s HSD post-hoc test on the CLR-transformed blood cell type data.

**Supplementary Table 3** *Plasmodium* stage proportion estimates calculated by CIBERSORT [21].

**Supplementary Table 4** Disease status reassignment after assessing the number of reads mapping to *Plasmodium* species.

**Supplementary Table 5** Total number of reads mapping to each *Plasmodium* gene for each sample.

**Supplementary Table 6** P-values from ANOVA tests between human gene expression levels and each known covariate.

**Supplementary Table 7** Enriched Reactome pathways for DE genes in *P. falciparum* and *P. vivax*-infected patients, along with Reactome pathways for DE genes in *P. falciparum* and *P. vivax*-infected patients which are unique to the Indonesian population.

**Supplementary Table 8** List of shared genes between *P. falciparum* and *P. vivax*-infected patients, along with a list of genes which are shared between *P. falciparum* and *P. vivax*-infected patients from Indonesia, Africa, and South America.

**Supplementary Table 9** List of all significant genes at an absolute log_2_ fold change threshold of one and an adjusted p-value of 0.05 for *Plasmodium*, as well as absolute log_2_ fold change and p-values for genes unique to Indonesia within individuals infected with *P. falciparum* or *P. vivax*.

**Supplementary Table 10** Gene Ontology analysis (Biological Processes) enrichment results.

**Supplementary Figure 1.**
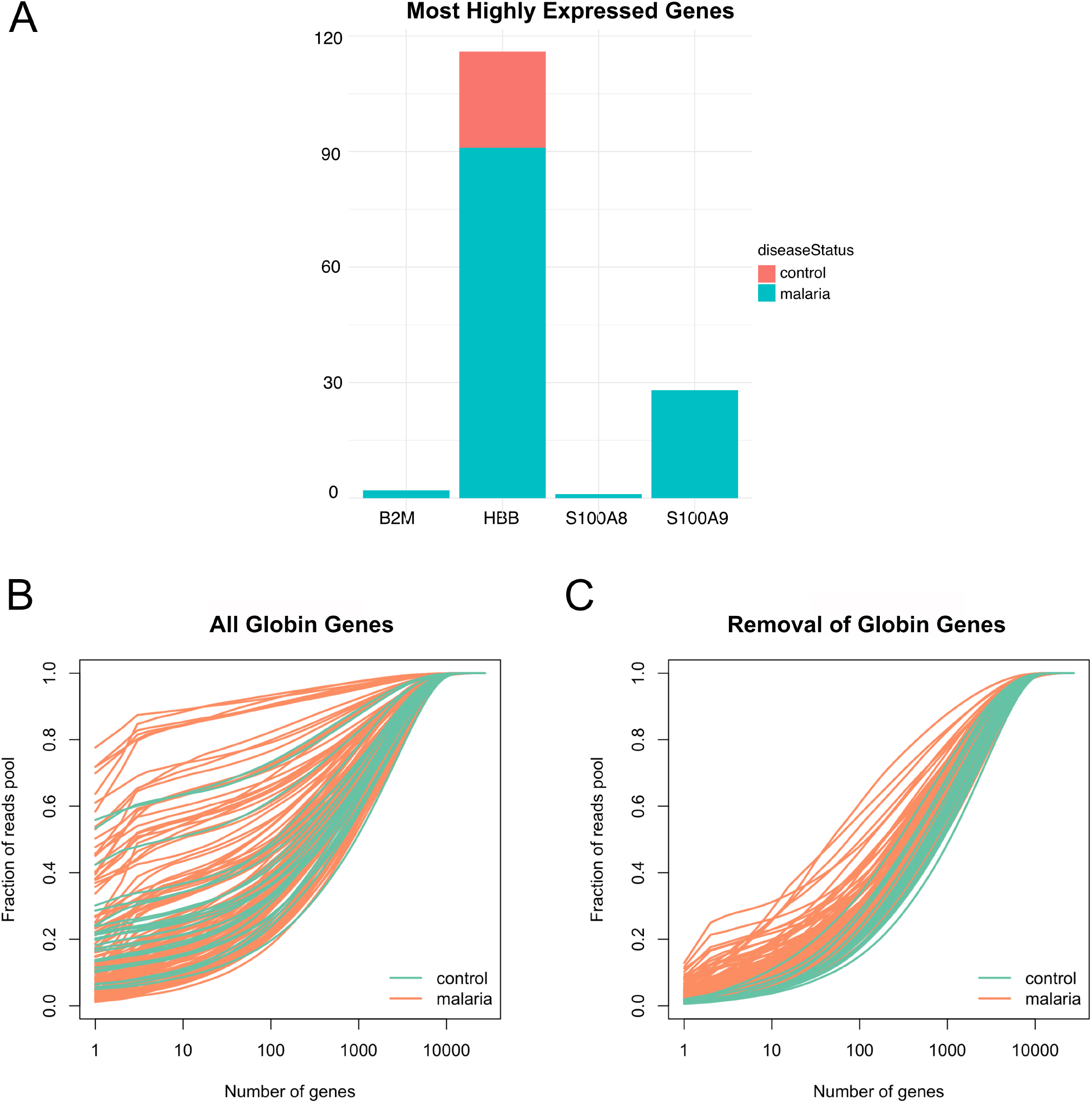
Globin gene removal in Yamagishi dataset. A) Most highly expressed genes in the Yamagishi transcriptome blood before removing globin genes. B) Rarefaction curves of the fraction of mRNAs contributed by the number of expressed genes within the Yamagishi dataset before filtering out globin genes. C) Rarefaction curves of the fraction of mRNAs contributed by the number of expressed genes within the Yamagishi dataset after filtering out globin genes. Orange lines indicate samples designated as having malaria by the study authors and green lines indicate samples being classified as healthy controls by the study authors.

**Supplementary Figure 2.**
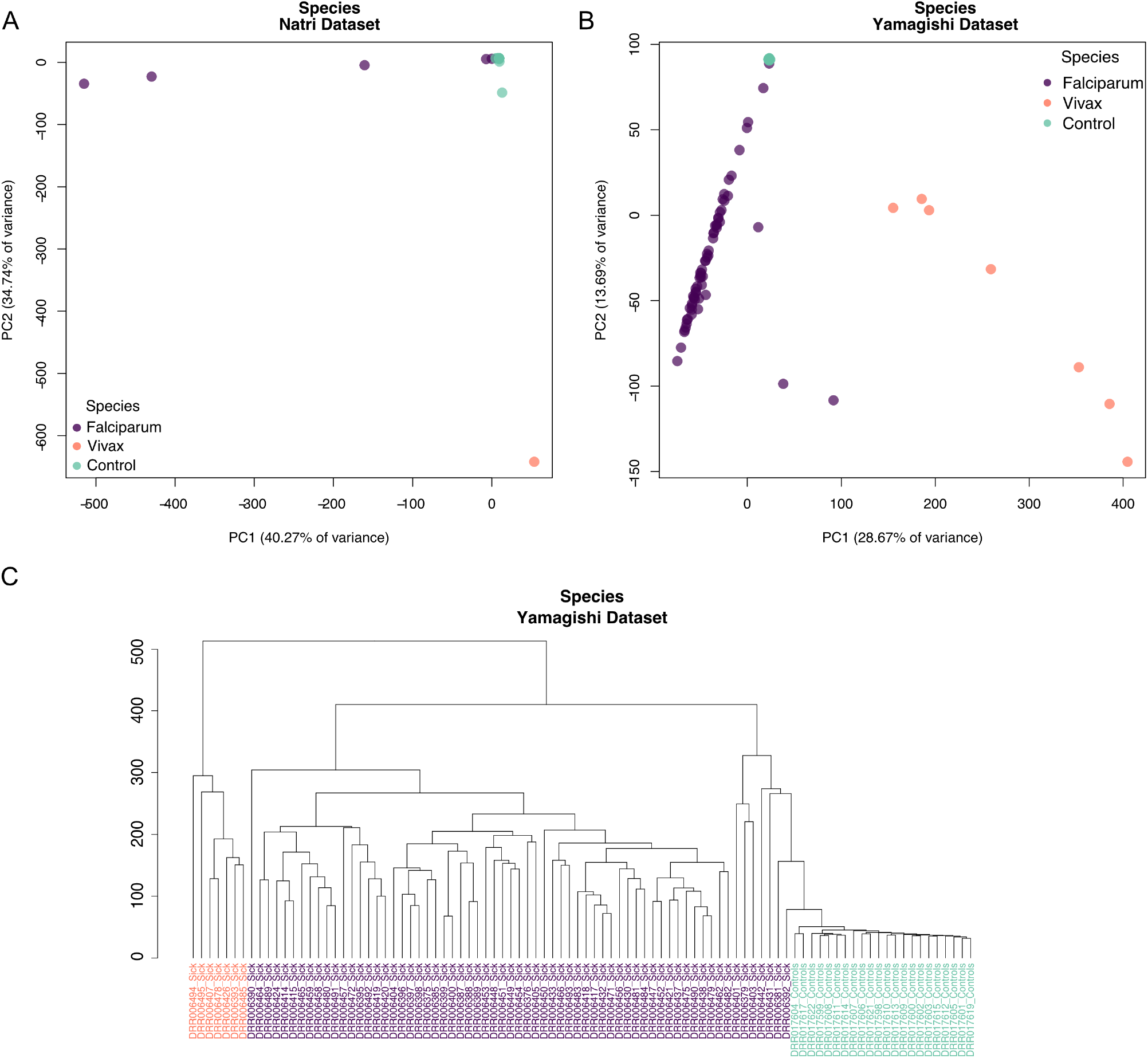
Species clustering analysis. A) Principal component analysis of reads mapping to *P. falciparum* and *P. vivax* within the Natri dataset. B) Principal component analysis of reads mapping to *P. falciparum* and *P. vivax* within the Yamagishi dataset. C) Hierarchical clustering by Euclidean distance of reads mapping to *P. falciparum* and *P. vivax* within the Yamagishi dataset. Controls are shown in green, *P. falciparum* in purple, and *P. vivax* in orange.

**Supplementary Figure 3.**
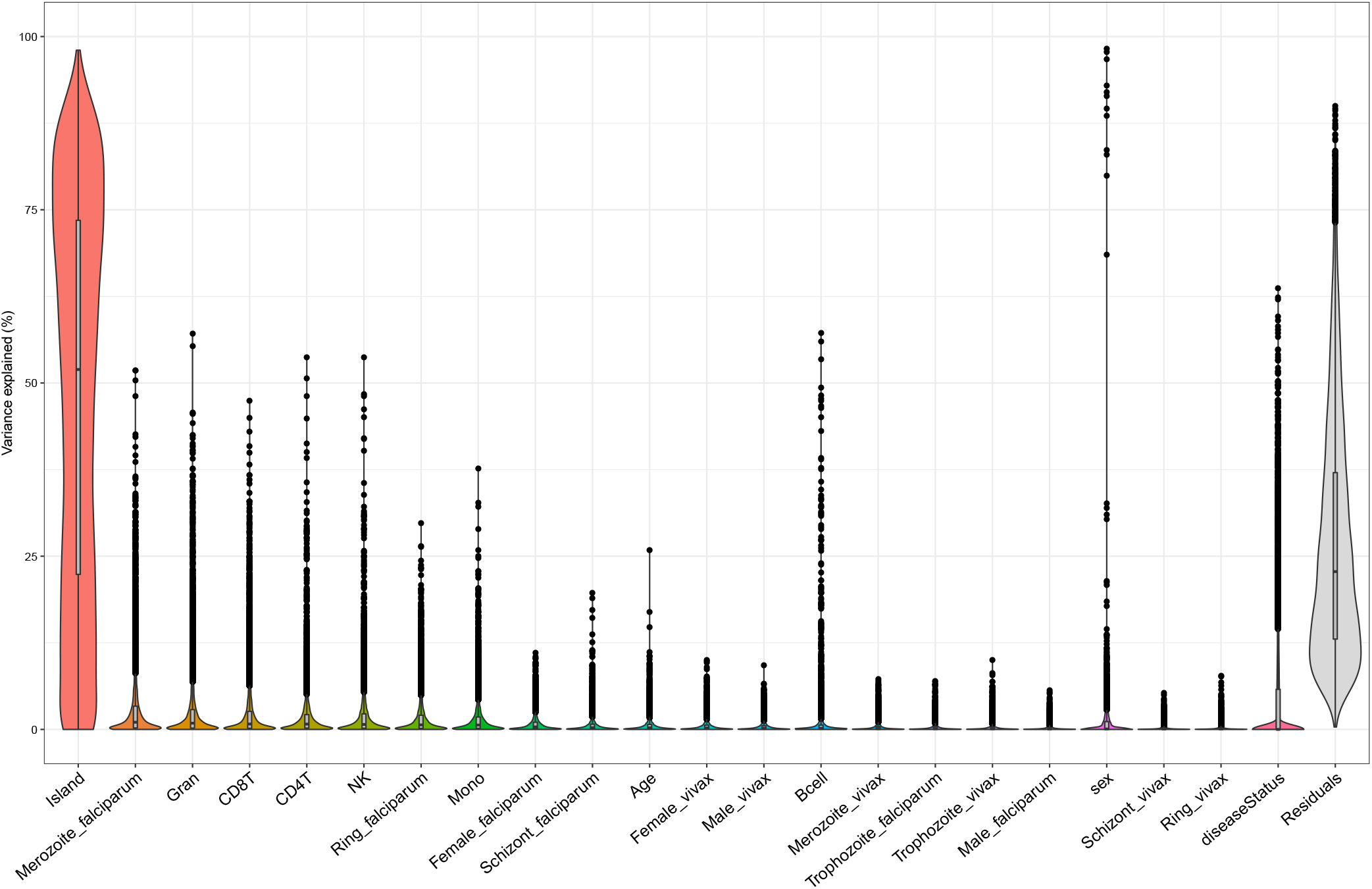
Percentage of variance explained by each covariate within the study design. Violin plots of the percentage of variance explained by all covariates used in the linear model. Estimates of variance explained were conducted by the R package variancePartition.

**Supplementary Figure 4.**
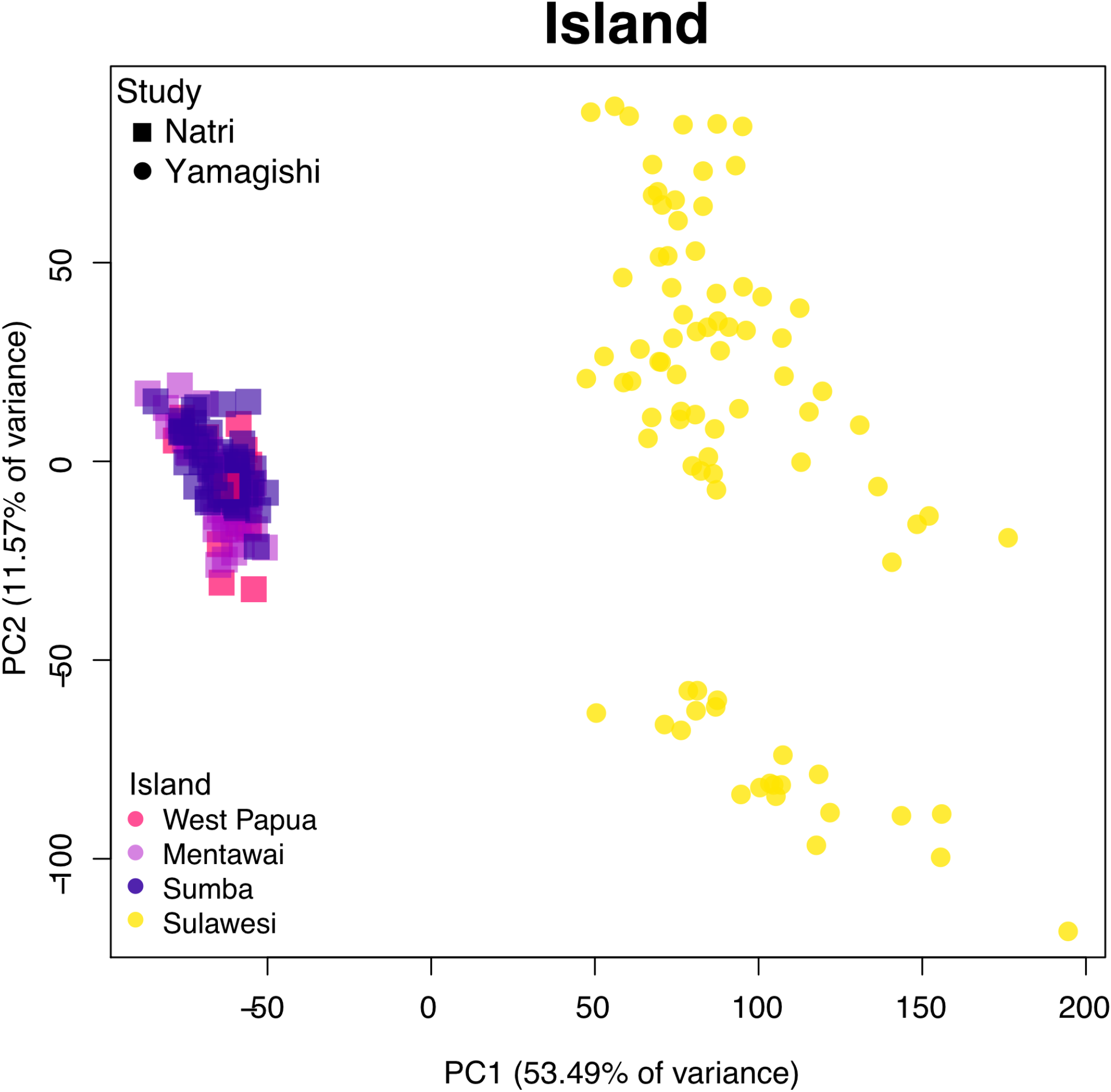
PCA analysis of island-level variation before batch correction.

**Supplementary Figure 5.**
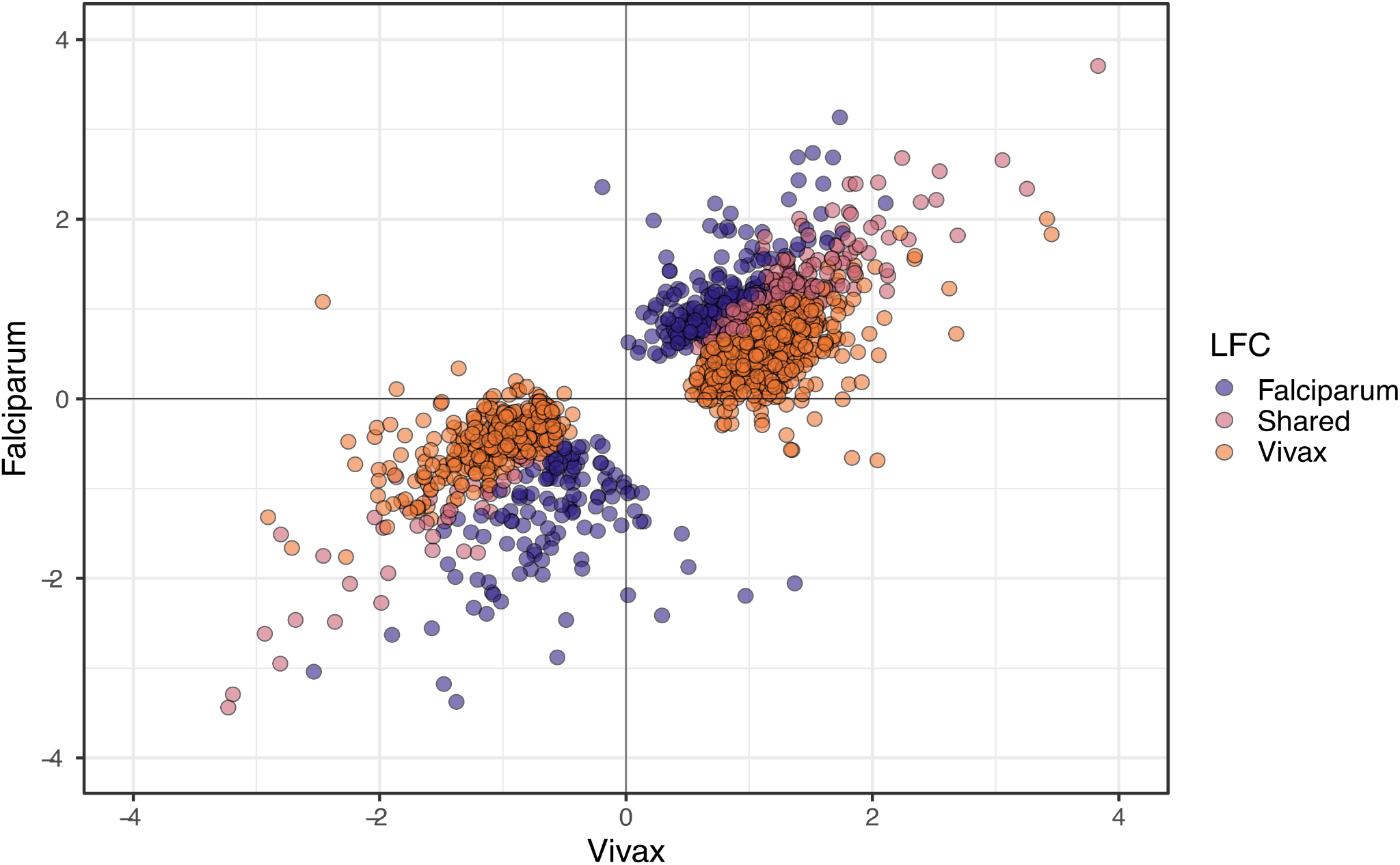
Within-gene variance in Indonesian malaria patients. log_2_ FC values of significant genes in *P. falciparum*-infected individuals, *P. vivax*-infected individuals, and genes shared between *P. falciparum* and *P. vivax*-infected individuals.

**Supplementary Figure 6.**
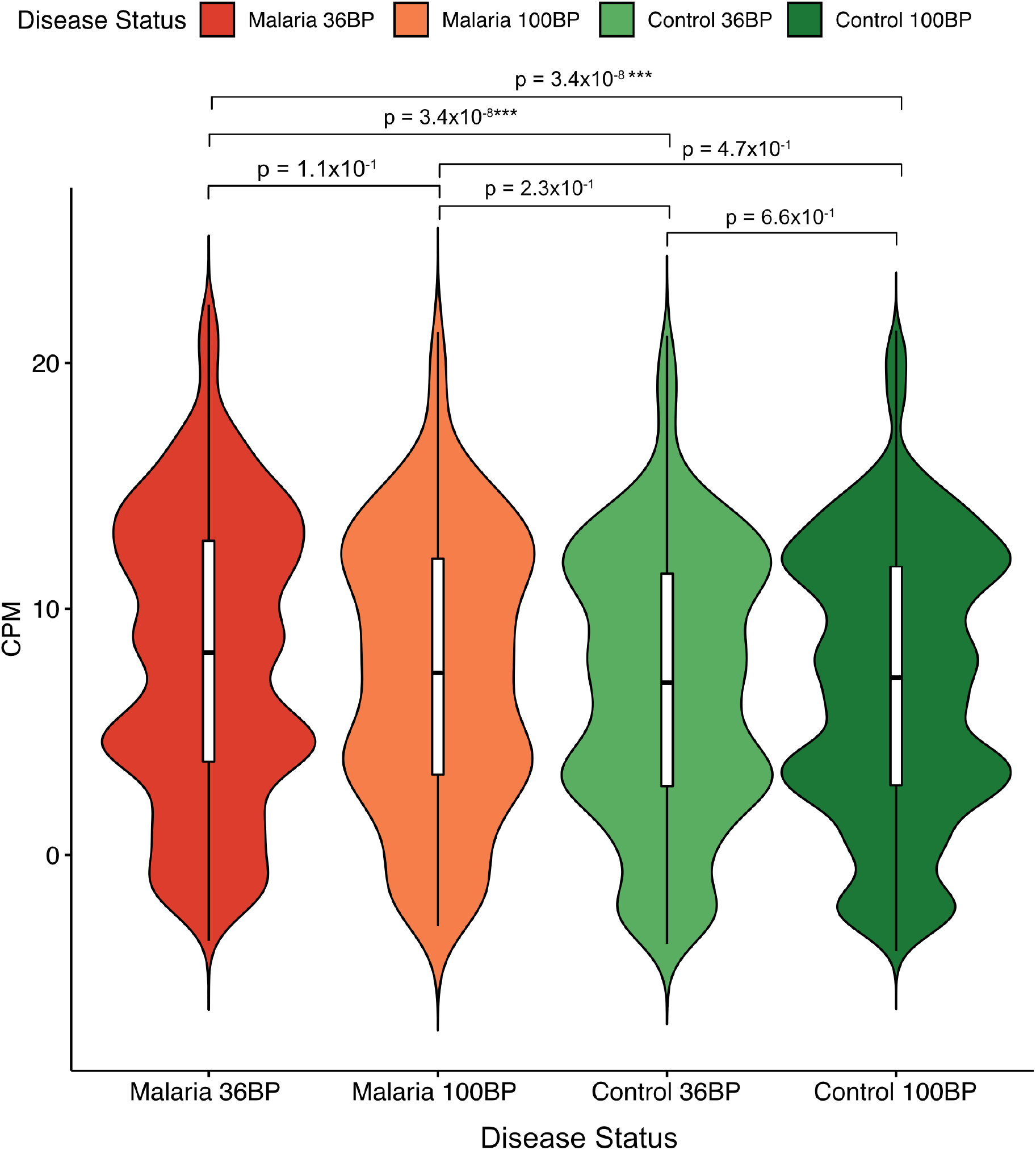
Enriched ribosomal protein genes by read length. Batch-corrected log_2_ CPM values of ribosomal protein genes (genes prefixed with “RPL”, “MRPL”, and RPS”) within cases of 36-BP read length, cases of 100-BP read length, and controls of 36 and 100-BP read length. Adjusted p-values from Tukey’s HSD are shown for group comparisons between cases of 36-BP length.

