## Supplementary figures and images for "Transcriptomic profiles of *Plasmodium falciparum* and *Plasmodium vivax*-infected individuals in Indonesia"

### Supplementary Figure 1

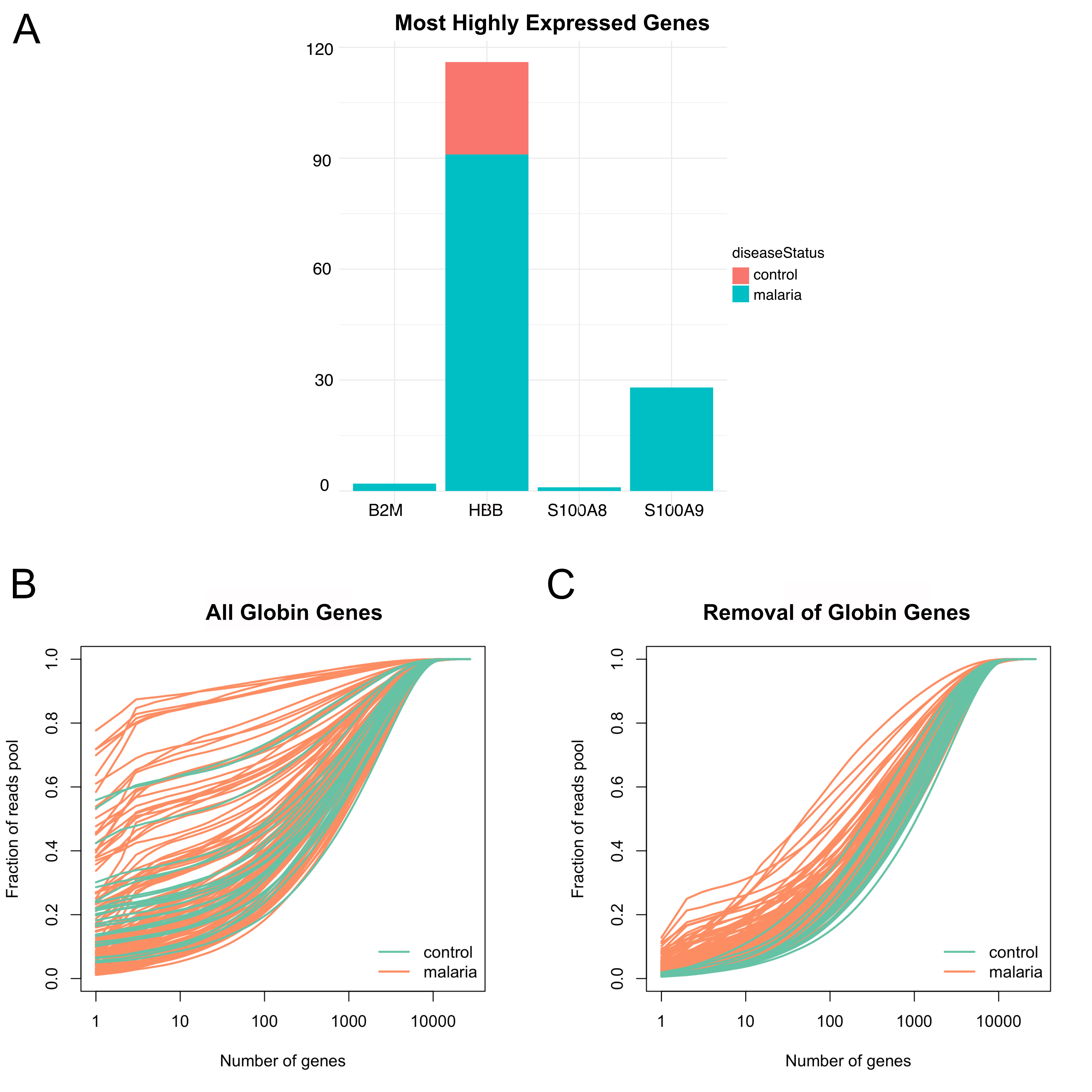

### Supplementary Figure 2

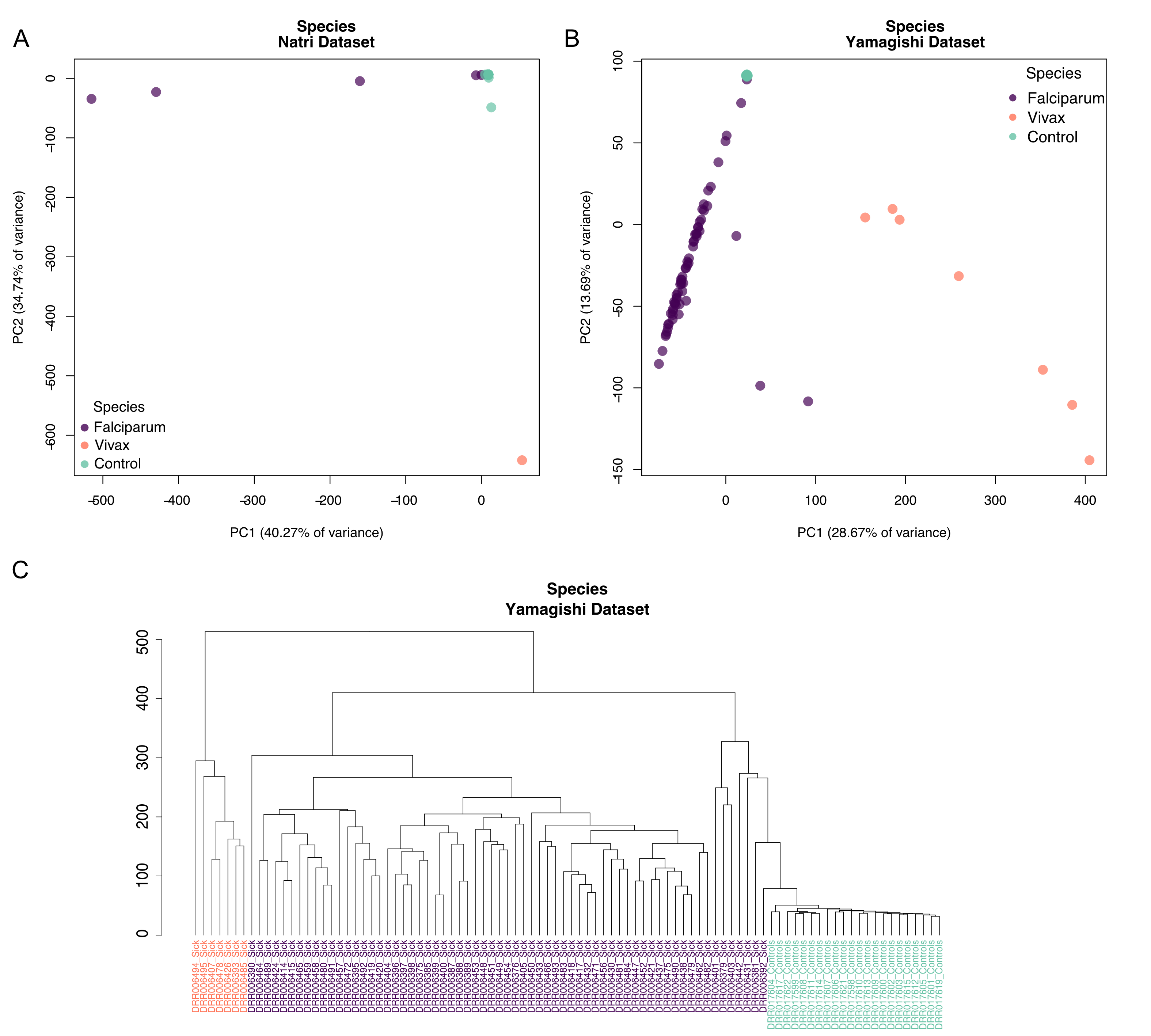

### Supplementary Figure 3

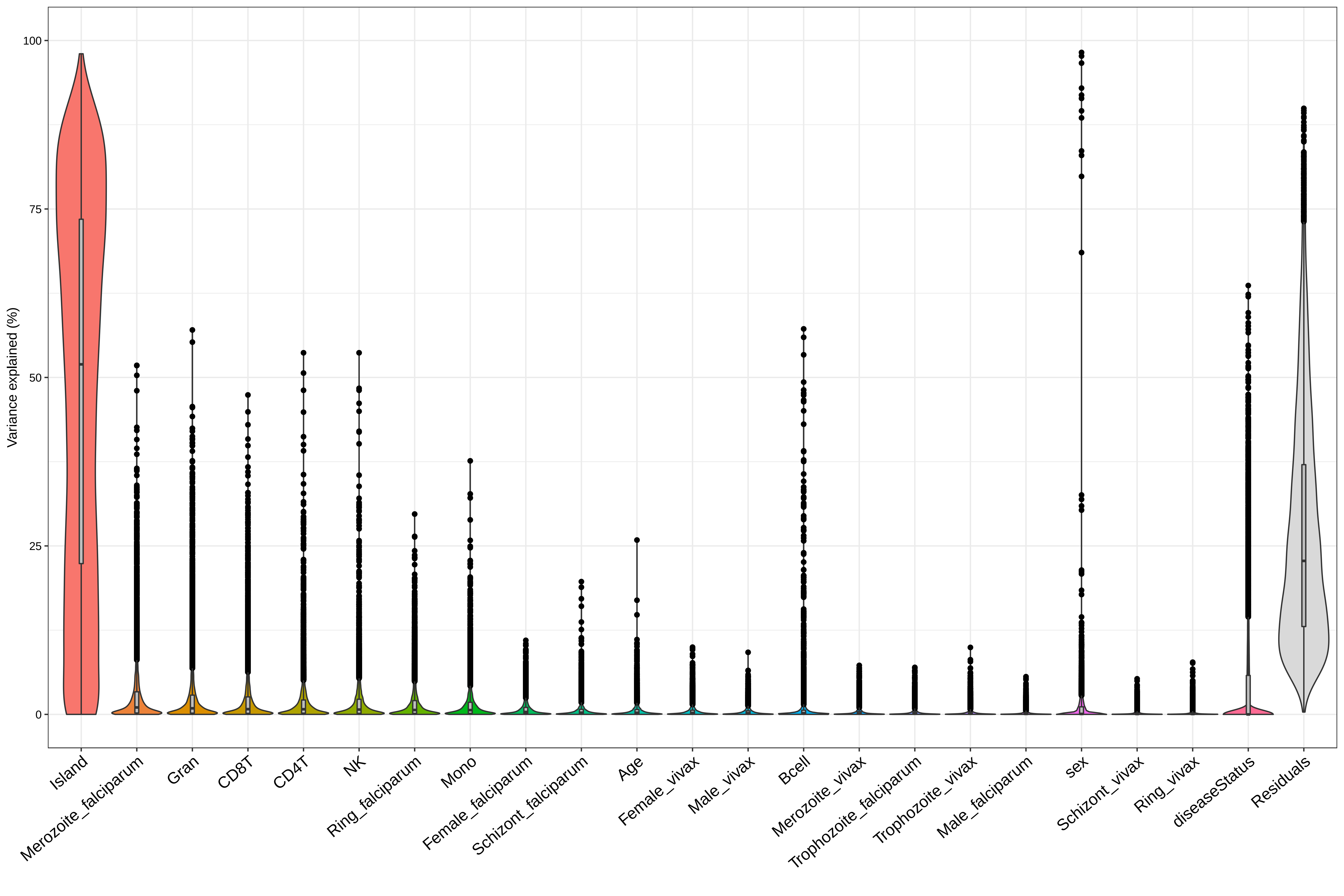

### Supplementary Figure 4

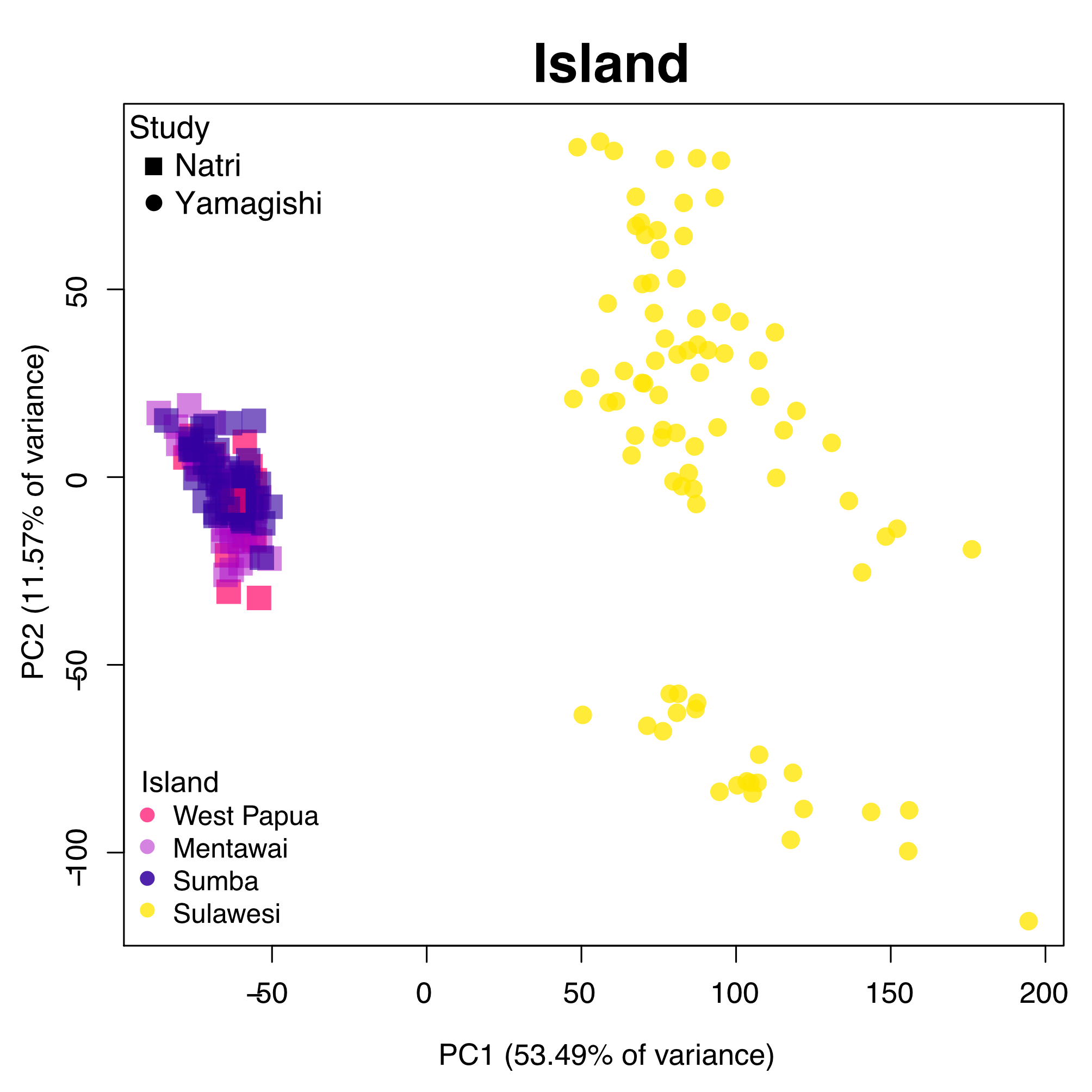

### Supplementary Figure 5

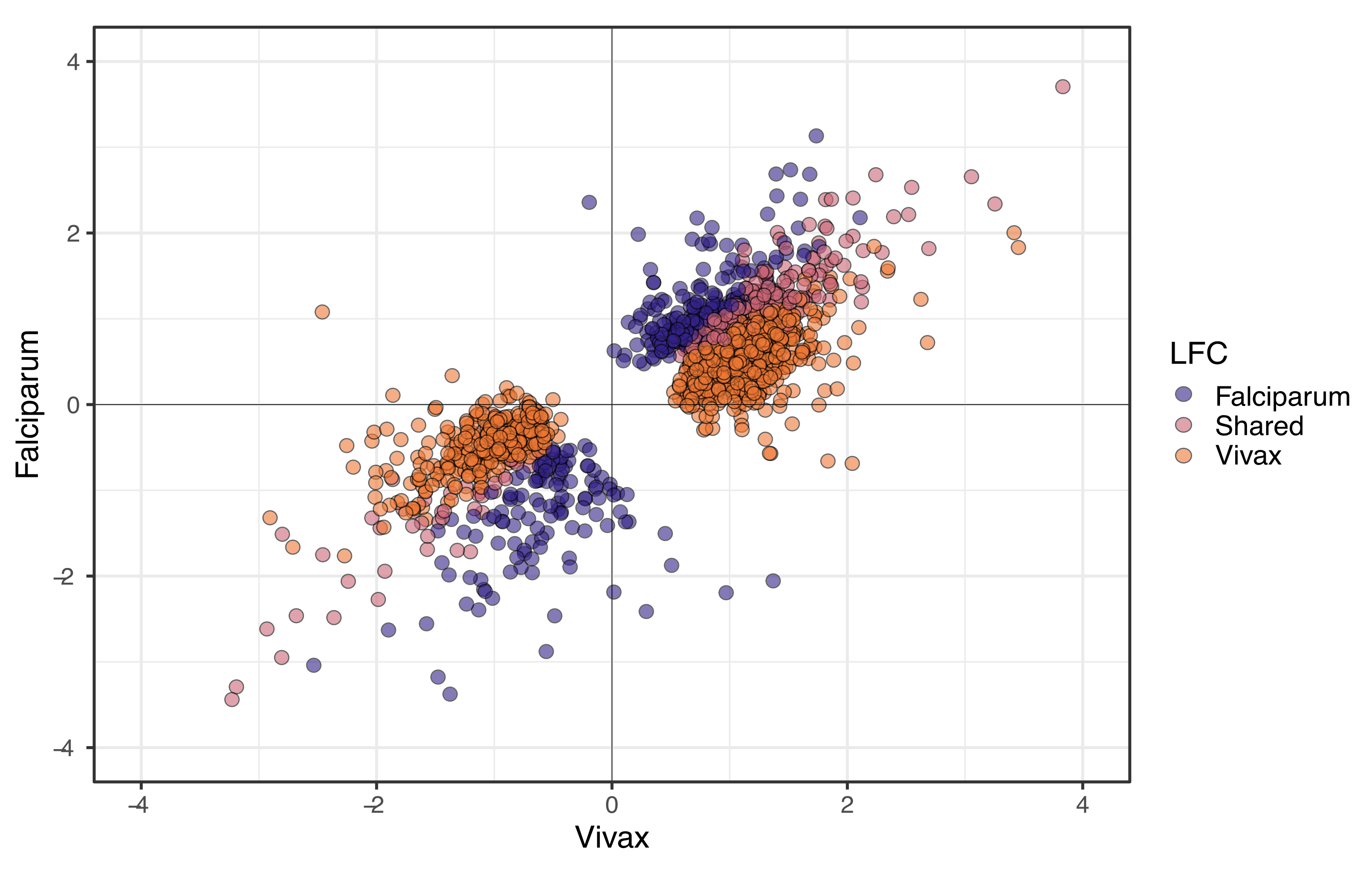

### Supplementary Figure 6

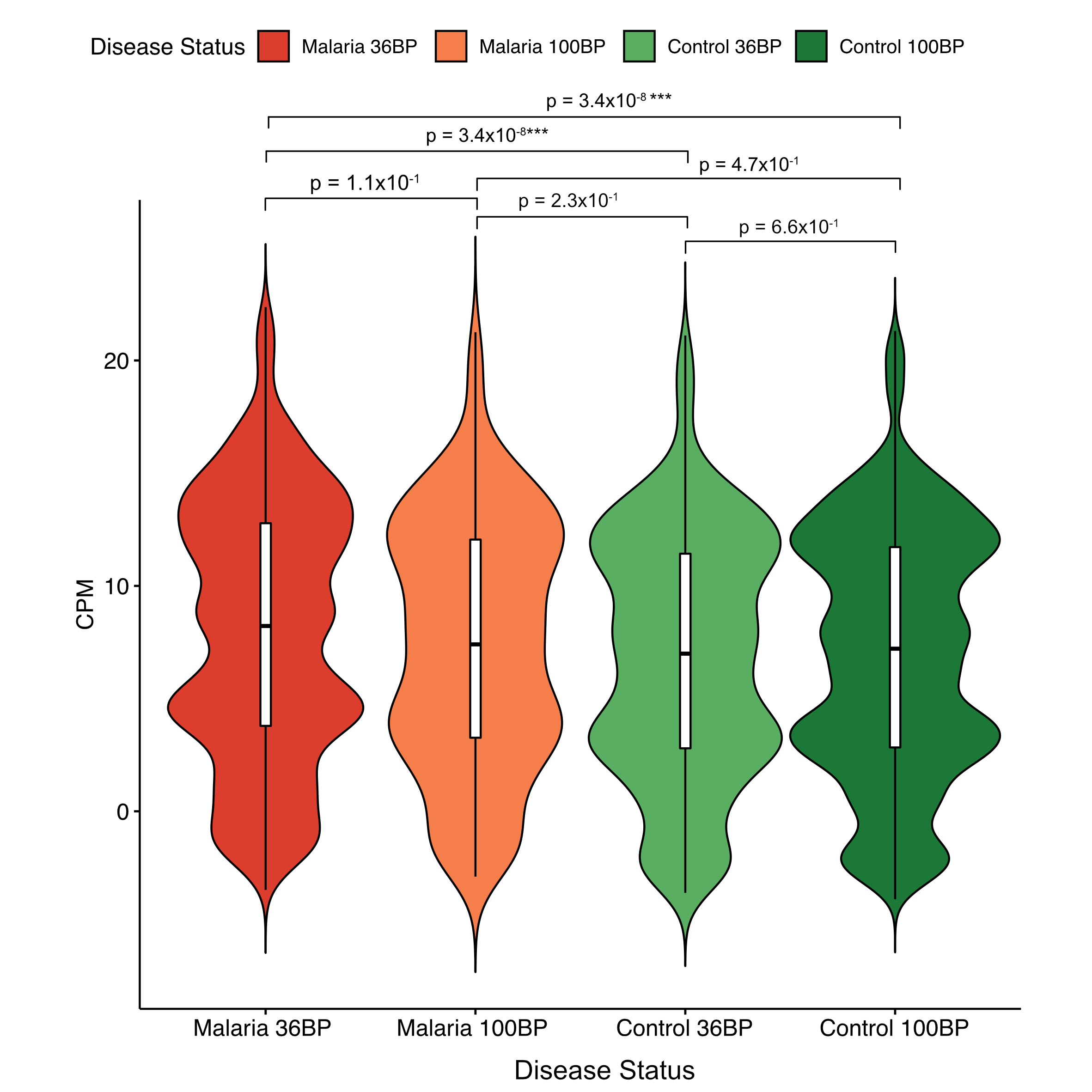
